# Alpha1-Adrenergic Receptor Mediated Long-Term Depression at CA3-CA1 Synapses Can be Induced via Accumulation of Endogenous Norepinephrine and is Preserved Following Noradrenergic Denervation

**DOI:** 10.1101/648873

**Authors:** Katie Dyer-Reaves, Anthoni M. Goodman, Amy R. Nelson, Lori L. McMahon

## Abstract

Locus coeruleus (LC) provides the sole source of noradrenergic (NA) innervation to hippocampus, and it undergoes significant degeneration early in Alzheimer’s disease (AD). Norepinephrine (NE) modulates synaptic transmission and plasticity at hippocampal synapses which likely contributes to hippocampus dependent learning and memory. We previously reported that pharmacological activation of α_1_ adrenergic receptors (α_1_ARs) induces long-term depression (LTD) at CA3-CA1 synapses. Here we investigated whether accumulation of endogenous NE via pharmacological blockade of NE transporters (NET) and the NE degradative enzyme, monoamine oxidase (MAO), can induce α_1_AR LTD, as these inhibitors are used clinically. Further, we sought to determine how degeneration of hippocampal NA innervation, as occurs in AD, impacts α_1_AR function and α_1_AR LTD. Bath application of NET and MAO inhibitors in slices from control rats reliably induced α_1_AR LTD when β adrenergic receptors were inhibited. To induce degeneration of LC-NA innervation, rats were treated with the specific NA neurotoxin DSP-4 and recordings performed 1-3 weeks later when NA axon degeneration had stabilized. Even with 85% loss of hippocampal NA innervation, α_1_AR LTD was successfully induced using either the α_1_AR agonist phenylephrine or the combined NET and MAO inhibitors, and importantly, the LTD magnitude was not different from saline-treated control. These data suggest that despite significant decreases in NA input to hippocampus, the mechanisms necessary for the induction of α_1_AR LTD remain functional. Furthermore, we posit that α_1_AR activation could be a viable therapeutic target for pharmacological intervention in AD and other diseases involving malfunctions of NA neurotransmission.

## Introduction

Noradrenergic (NA) input from the locus coeruleus (LC) to hippocampus is critical for hippocampus-dependent learning and memory (Gibbs et al., 2010; Harro et al., 1999; Koob et al., 1978; Lemon et al., 2009), and its degeneration in Alzheimer’s disease (AD) has been well documented (Forno, 1966; Yamada and Mehraein, 1977; Zarow et al., 2003). Specifically, loss of LC cell bodies positively correlates with the severity of dementia and duration of the disease in AD patients. In fact, the LC is the first target of pretangle tau pathology, with measurable cell loss occurring in the prodromal phase of AD (Arendt et al., 2015; Braak et al., 2011; Chalermpalanupap et al., 2017; Grudzien et al., 2007; Jucker and Walker, 2011) that may mediate the emergence of cognitive changes (Arendt et al., 2015; Braak et al., 2011; Chalermpalanupap et al., 2017; Kelly et al., 2017; Theofilas et al., 2017). Despite this body of evidence, the effect of early LC degeneration on hippocampal synaptic function is poorly understood.

The LC is the sole provider of NA innervation in hippocampus (Aston-Jones, 2004), and norepinephrine (NE) modulates synaptic efficiency critical for learning and memory (Hagena et al., 2016; Harley, 1991, 2007; Kemp and Manahan-Vaughan, 2008; Scheiderer et al., 2004). Selective electrolytic lesioning using the neurotoxin N-(2-chloroethyl)-N-ethyl-2-bromobenzylamine (DSP4) or silencing of LC neurons using optogenetics in murine models leads to deficits in learning, memory, and cognitive flexibility (Harro et al., 1999; Janitzky et al., 2015; Koob et al., 1978). NE, via activation of adrenergic receptors (ARs), modifies the strength of synaptic transmission at glutamatergic synapses and the ability of these synapses to undergo long-term plasticity (Bramham et al., 1997; Bröcher et al., 1992; Erickson et al., 1997; Harley, 1991; Harley and Sara, 1992; Hopkins and Johnston, 1984; Izumi and Zorumski, 1999; Katsuki et al., 1997; Thomas and Palmiter, 1997a, 1997b, 1997c). Accordingly, blockade of β adrenergic receptors (ARs) reduces NMDAR-dependent LTP in hippocampal slices (Harley, 1991), while NE application facilitates the induction of LTP in dentate and spatial memory formation through activation of β-ARs (André et al., 2015; Bröcher et al., 1992; Hopkins and Johnston, 1984; Izumi and Zorumski, 1999; Katsuki et al., 1997; Kemp and Manahan-Vaughan, 2008). Along with β-ARs, α-ARs are necessary for spatial memory learning tasks, as α_1_AR agonists enhance and antagonists block the formation of memory (Pussinen et al., 1997; Puumala et al., 1998). Activation of α_1_- and β-ARs by NE can also facilitate tetanus-induced LTP at mossy-fibers synapses in area CA3 (Hopkins and Johnston, 1984; Huang et al., 1996). Transgenic mice harboring constitutively active α_1_ARs have enhanced learning and memory, while α_1_AR knock-out mice showed deficits compared to WT (Collette et al., 2014; Doze et al., 2011). We previously reported that NE, or the selective α_1_AR agonist methoxamine, induces a form of long-term depression (LTD) at CA3-CA1 synapses that is dependent on NMDARs, Src kinase, and extracellular signal-regulated protein kinase (ERK) activation (Scheiderer et al., 2004, 2008).

Given that activation of α_1_ARs using exogenous agonists induces LTD at hippocampal CA3-CA1 synapses (Scheiderer et al., 2004, 2008), we wanted to determine if increasing endogenous extracellular NE accumulation via pharmacological inhibition of the norepinephrine transporter (NET) and the degradative enzyme monoamine oxidase (MAO) similarly induces α_1_AR LTD. This is important since these inhibitors are widely used as therapeutic treatments in disorders such as ADHD and depression, where imbalances in catecholamine neurotransmission, specifically NE, are known to occur (Castellanos et al., 1996; Israel, 2015; Vanicek et al., 2014; Zametkin and Rapoport, 1987). Furthermore, because LC degeneration, and loss of hippocampal NA innervation, is clinically relevant to normal aging, AD, and Parkinson’s disease (PD) (Mann, 1983; Mann et al., 1983; Marien et al., 2004; Szot, 2012), we set out to investigate the impact of NA degeneration on the ability of pharmacological activation of α_1_ARs to induce LTD at hippocampal CA3-CA1 synapses.

## Materials and methods

### Animal care

All experiments were conducted with an approved protocol from the University of Alabama at Birmingham Institutional Animal Care and Use Committee in compliance with the National Institutes of Health guidelines. All efforts were made to minimize animal suffering and to reduce the number of animals used. Six-week old male Sprague Dawley rats (Charles River) were used in all experiments. Animals were housed two per cage and were kept on a 12-hour light/dark cycle with ad libitum food and water.

### LC lesion

Hippocampal NA denervation was performed using the NA axon specific neurotoxin DSP-4 (Tocris, Ellisville, MO), known to cause terminal retrograde degeneration by targeting the NE uptake system (Jaim-Etcheverry and Zieher, 1980; Ross and Stenfors, 2014). DSP-4, delivered intraperitoneally (IP), readily crosses the blood-brain-barrier and targets NA axons of the central nervous system while peripheral NA systems are unaffected (Fritschy and Grzanna, 1989; Jaim-Etcheverry and Zieher, 1980; Ross and Stenfors, 2014). NE uptake in these axons is rapidly blocked with maximum effect achieved by 4-6 hours and within 4-5 days DβH immunoreactivity declines (Fritschy et al., 1990; Ross, 1976). NA sprouting occurs in a region-specific manner over approximately 5 weeks (Booze et al., 1988) along with an observed 57% decline in LC cell bodies one year after DSP-4 treatment (Fritschy and Grzanna, 1992). Rats were lightly anesthetized with isofluorane and injected IP with DSP-4 (50mg/kg) in saline or saline alone at 48-hour intervals for a total of 3 injections.

### Drugs and solutions

All drugs (Sigma, St. Louis, MO) were prepared as stock solutions and diluted to the appropriate working concentration at the time of electrophysiology. Phenylephrine (Phe, α_1_AR agonist; in deionized water), propranolol (β-AR antagonist; in DMSO) and prazosin (α_1_AR antagonist; in DMSO) were prepared fresh daily and atomoxetine (Atmx, NET inhibitor; in deionized water) and clorgiline (Clor, MAO inhibitor; in deionized water) were frozen in 300μL aliquots until used for recordings.

### Slice preparation and electrophysiological recordings

α_1_AR LTD was induced using the α_1_AR agonist phenylephrine (Phe,100μM) that was bath applied for 10 or 15 min following a 20 min stable baseline of recorded fEPSPs as done previously (Scheiderer et al., 2004, 2008). Experiments to test whether extracellular accumulation of endogenous NE could induce LTD, as well as to test the functionality of the NA fibers remaining following DSP-4-mediated lesion, slices were exposed to bath application of the NET inhibitor Atmx (500nM) plus the MAO inhibitor Clor (10μM) for 10 or 20 min following stable baseline transmission. Previous reports have documented the ability of NET inhibition to block reuptake of NE and induce increases in extracellular NE (Youdim and Riederer, 1993). Additionally, selective inhibition of MAO, the enzyme responsible for NE degradation, has also been shown to cause accumulation of NE extrasynaptically (Youdim and Riederer, 1993). Because NE has similar affinity for α- and β-ARs, it was necessary to pharmacology inhibit β-AR activation with propranolol (10µm) to reveal LTD following accumulation of extracellular NE. To ensure the α_1_AR specificity of the LTD induced by endogenous NE in these recordings, interleaved experiments were conducted in the presence of 10μM prazosin (in addition to the cocktail of Clor, Atmx and propranolol, which is referred to as CAP).

### Immunohistochemistry

Following electrophysiological recordings, 400μm-thick hippocampal slices were stored in 4% paraformaldehyde at 4°C until the time of staining. Twenty-four hours prior to staining, slices were rinsed in phosphate buffered saline (PBS) and then transferred to a 30% sucrose/PBS solution. Tissue was resectioned to 50μm using a freezing microtome. Sections were washed 3 x 10 min in PBS at room temperature and incubated in blocking buffer (10% normal donkey serum (NDS) in 0.3% PBS Triton/PBS) for 90 minutes. Primary antibodies [rabbit anti-tyrosine hydroxylase (TH, 1:200) and mouse anti-dopamine β-hydroxylase (DβH, 1:300); Chemicon, Temecula, CA] were diluted in blocking buffer, and applied to free-floating sections and incubated overnight at 4°C. Slices were washed 3 x 10 min with PBS and were labeled with fluorescence-activated secondary antibodies [donkey anti-rabbit Alexa 594 (1:200) and donkey anti-mouse Alexa 488 (1:200); Invitrogen, Eugene, OR] diluted in blocking buffer and incubated for 1-2 hours at room temperature. Slices were washed 3 x 30 min and incubated with Hoescht nuclear stain [1μl stock/10ml PBS] for 15 min at room temperature. Slices were mounted on slides using Permafluor (Immunon, Waltman, MA) and viewed on a Leica (Exton, PA) DM IRBE laser scanning confocal microscope. The CA1 s. radiatum within the field of view from one slice per animal was analyzed using ImageJ software. Sequential scans of blue, green, and red channels were obtained and ~20μm stacks of images were collected in a z-axis of 1.0-1.5μm step size, averaging 2 scans per image. Maximum projections were generated and used for NE fiber quantification. DβH-positive fibers were measured and counted, following the criterion that only fibers with 4 or more consecutive boutons be considered as a fragment of axon.

### Data analysis

Electrophysiology data were analyzed using custom-written Labview data acquisition and analysis software (Scheiderer et al., 2004, 2008) after being filtered at 3 kHz and digitized at 10 kHz. The fEPSP slope was measured and evaluated as a series of 5 averaged raw data points plotted versus time. The LTD magnitude was calculated by comparing the average fEPSP slope recorded during the last 10 min of baseline transmission to the slope at 20 min post drug washout. When more than one slice was recorded per animal for a given experiment (e.g. Phe or Amtx+Clor), the data were averaged together to represent the finding for that animal. Therefore, the reported *n* refers to animal number. Paired student’s *t-*tests were used for statistical analysis within groups. Unpaired student’s *t-*tests or one-way ANOVAs were used to evaluate statistical significance between groups. The significance level was set at p<0.05 and the data are presented as the mean ± s.e.m.

## Results

### α_1_AR activation induces LTD at CA3-CA1 synapses in rat hippocampus

Our laboratory previously reported that bath application of NE or the selective α_1_AR agonist methoxamine is sufficient to induce a NMDAR-dependent LTD of extracellular fEPSPs at CA3-CA1 glutamatergic synapses in hippocampal slices (Scheiderer et al., 2004, 2008). Here, we show that application of another α_1_AR agonist, Phe (100μM; phenylephrine) also reliably induces α_1_AR LTD (Fig. 1A, 84 ± 4% of baseline fEPSP slope (n=6); p<0.01) similar to our previous report (Scheiderer et al., 2004, 2008). To test whether α_1_AR LTD can also be induced via extracellular accumulation of endogenous NE in hippocampus, the selective NET inhibitor Atmx (500nM) was applied together with an inhibitor of NE degradative enzyme MAO, Clor (10μM). Bath application of to Atmx and Clor did not elicit a significant change in synaptic strength compared to baseline (Fig. 1B1, 94 ± 5% of baselined fEPSP slope, n=6, p=0.14). However, when we evaluated individual experiments there were clear cases of Atmx and Clor induced depression (Fig. 1B2) and potentiation (Fig. 1B3) of the extracellular fEPSP that were masked when all of the experiments were averaged. This variable response can be attributed to coincident global activation of α_1_, α_2_, and β-ARs, as all of these receptors are located at pre- or post-synaptic locations at CA3-CA1 synapses.

**Figure 1.**
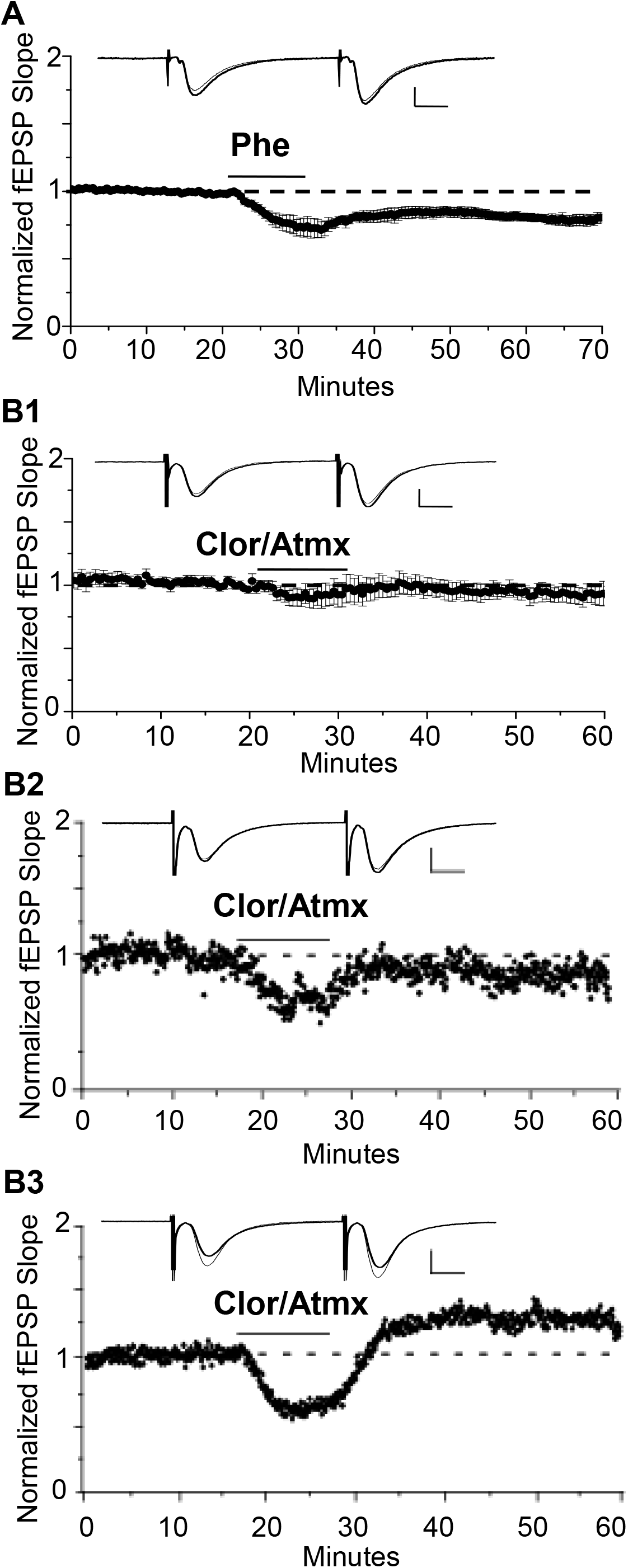
LTD is induced by α_1_AR agonist, phenylephrine, or endogenous NE. A. α_1_AR LTD is induced by the selective α_1_AR agonist Phe. 100 µM is able to induce α_1_AR LTD in control animals (84±4% of baselined fEPSP slope, n 6), (scale bar: 0.5mV, 10ms). B. Collective activation of α_1_-, α_2_, and β-ARs by endogenous NE results in variable effects on synaptic efficacy. B1. Averaged experiments in Clor/Atmx results in no significant change in baseline transmission (94±5% of baseline fEPSP slope, n=6). B2. Representative example of LTD induced via NET and MAO inhibition by Atmx (500nM) and Clor (10µM, respectively). B3. Single example of LTP induced via endogenous NE accumulation in the presence of Clor/Atmx.

Because activation of β-ARs causes potentiation at CA3-CA1 synapses (Bröcher et al., 1992; Hopkins and Johnston, 1984; Izumi and Zorumski, 1999; Katsuki et al., 1997) their activation by endogenous accumulation of NE could be masking possible α_1_AR LTD expression induced by Atmx and Clor application. To determine whether blockade of β-AR activation would unmask LTD, propranolol (10μM) was applied for the duration of the recording period during the Atmx and Clor experiments (collectively named CAP), and this resulted in reliable, and significant LTD expression (Fig. 2A, 83 ± 6% of baseline fEPSP slope, n=4, p=0.02). The LTD magnitude induced by the selective α_1_AR agonist Phe (Fig. 1A) is not significantly different from that induced by the combination of NET, MAO, and β-AR inhibition (Fig. 2A) (p=0.8). To confirm that the LTD following accumulation of endogenous NE is also mediated by α_1_AR activation, interleaved experiments were performed with the α_1_AR antagonist prazosin (10μM) resulting in a complete block of LTD (Fig. 2B, CAP plus prazosin, 96.5 ± 4% of baseline fEPSP slope, n=3 p=0.4).

**Figure 2.**
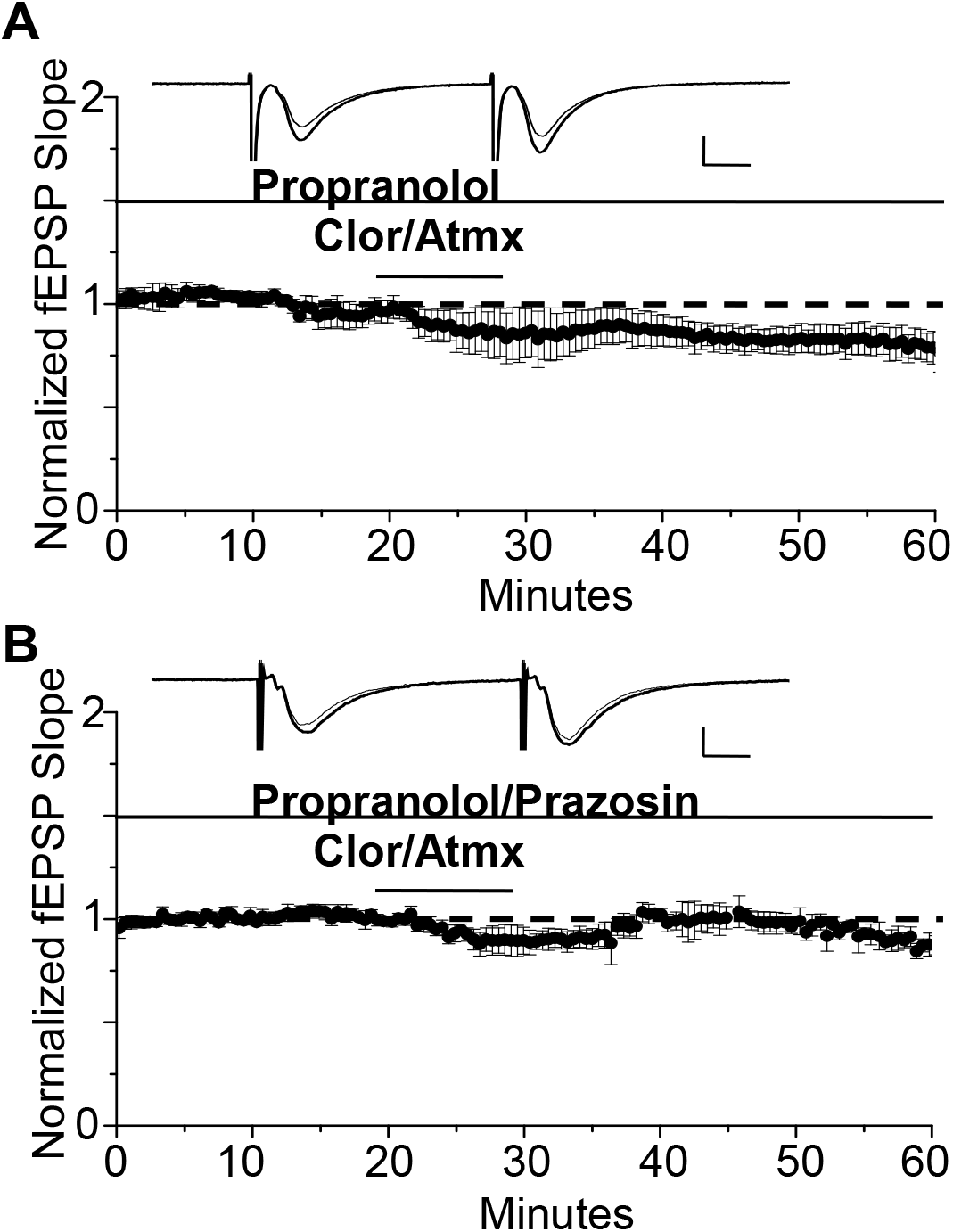
The β AR antagonist propranolol is able to unmask α_1_AR LTD when used in addition to Clor and Atmx. A. In the presence of the β AR antagonist propranolol (1µM) accumulation of endogenous NE induces α_1_AR LTD (83±6%of baselined fEPSP slope, n=4). B. α_1_AR LTD is prevented by application of the α_1_AR antagonist prazosin (10µM) in the presence of propranolol, Clor, and Atmx (p>0.05).

### DSP-4 causes a significant decrease in NA innervation in CA1 of hippocampus

To determine whether loss of NA input to hippocampus is sufficient to cause deficits in α_1_AR LTD, the NE specific neurotoxin DSP-4 (50mg/kg, in saline), was administered intraperitoneally at 48-hour intervals for a total of 3 injections (control animals received injections of saline only). DSP-4 targets the NE uptake system and induces alkylation of vital neuronal structures (Ross, 1976) causing degeneration of hippocampal NA innervation (Fritschy et al., 1990; Fritschy and Grzanna, 1989; Jonsson et al., 1981). This robust treatment protocol was used because a previous study reported that mice treated with one dose of DSP-4 had an increased probability of hippocampal NA axon sprouting compared to mice treated 3 times with the toxin (Puoliväli et al., 2000). NA innervation in s. radiatum of area CA1 following DSP-4 treatment was evaluated using anti-DβH immunohistochemical staining of NA fibers, which were then imaged via confocal microscopy. DSP-4 induced a significant decrease in NA fiber number (Fig. 3B) (*F*(3,19)=23.28, p<0.001) but no change in individual fiber length in CA1 s. radiatum in animals sacrificed 7-21 days following the first injection (Fig. 3C) (*F*(3,658)=2.03, p=0.108) This protocol led to a reduction of ~85% of DβH-positive immunostaining. Interestingly, the morphology of some of the remaining DβH-positive fibers in the DSP-4-treated animals have a thicker, knobby appearance compared to the thin, more delicate fibers found in the control group (Fig. 3A arrows).

**Figure 3.**
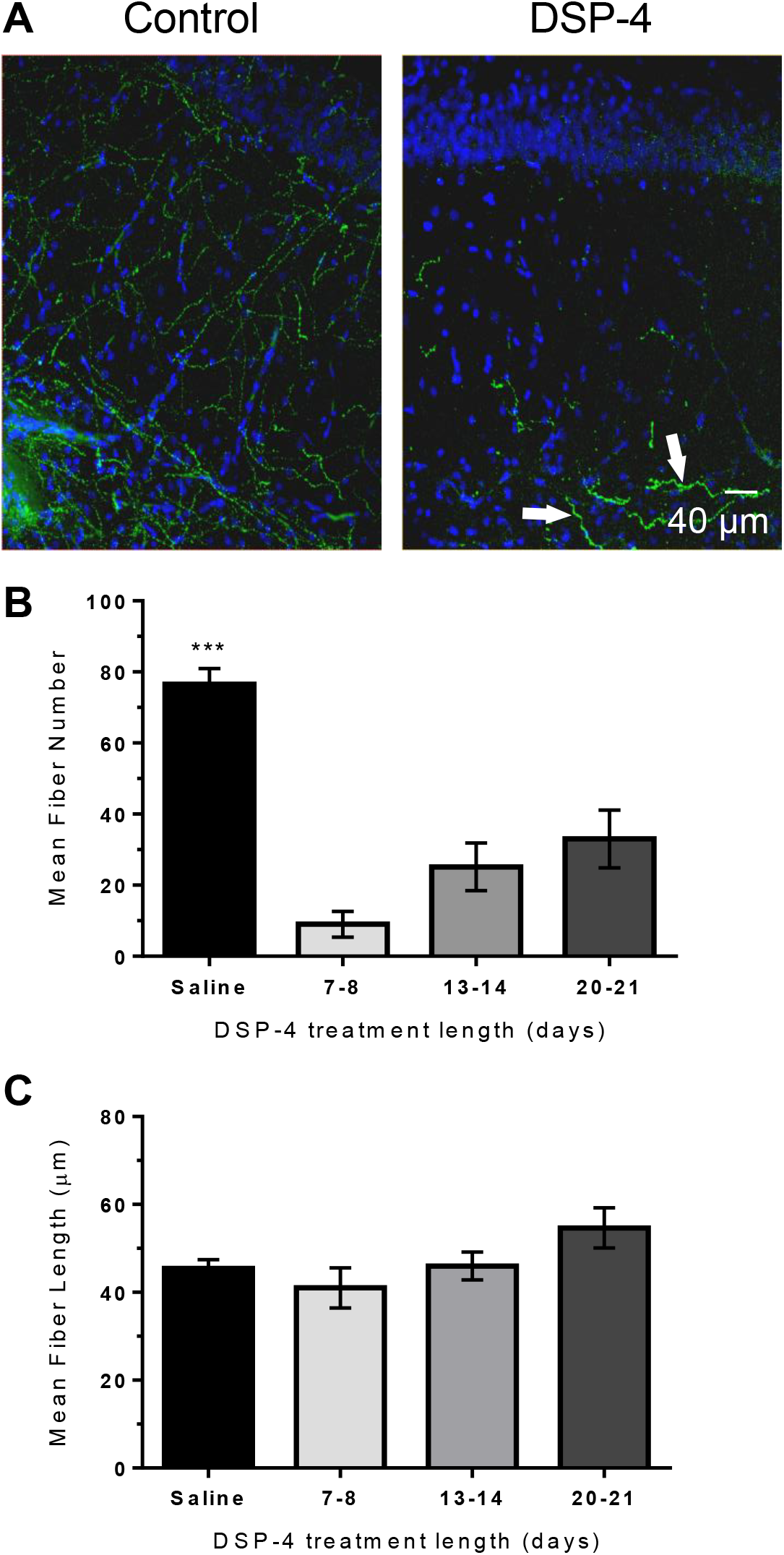
DSP-4 treatment causes significant NE degeneration in CA1. of hippocampus. A. Anti-DβH (green) and DAPI (blue) hippocampal staining of NE fibers in area CA1 s radiatum from a saline-treated (control) animal or following DSP-4 treatment (scale bar: 40µm). Arrows denote thicker, knobby appearance of some remaining NE fibers. B. Total fiber number in DSP-4 treated animals in CA1 s. radiatum is significantly decreased compared to saline-treated measured 7-21 days post treatment. C. Significant degeneration of NE fiber length results after DSP-4 treatment. C. Total NE fiber number is significantly decreased by DSP-4 treatment by ~85% (saline, n=7; DSP-4, n=25). D. Total fiber number in DSP-4 treated animals in CA1 s. radiatum is significantly decreased compared to saline-treated measured 7-21 days post treatment.

### α_1_AR LTD remains intact following NA-fiber degeneration

We next tested whether systemic treatment with DSP-4 and subsequent ~85% reduction of NA innervation would negatively impact the ability of direct activation of α_1_ARs by the selective α_1_AR agonist Phe to induce LTD. Surprisingly, we found that the magnitude of α_1_AR LTD was not significantly different between saline-treated and DSP-4 treated animals. (Fig. 4A, DSP-4: 85 ± 3%, n=7, p<0.001 vs Fig. 4B, saline v. DSP-4, p=0.85). Thus, despite an ~85% decrease in NA innervation in s. radiatum of area CA1, α_1_ARs remain coupled to downstream signaling cascades (Src kinase and pERK) (Scheiderer et al., 2008) necessary for the induction of LTD. However, it is unclear whether α_1_AR LTD can be induced by endogenously released NE from the remaining 15% of NE fibers following DSP-4 treatment.

**Figure 4.**
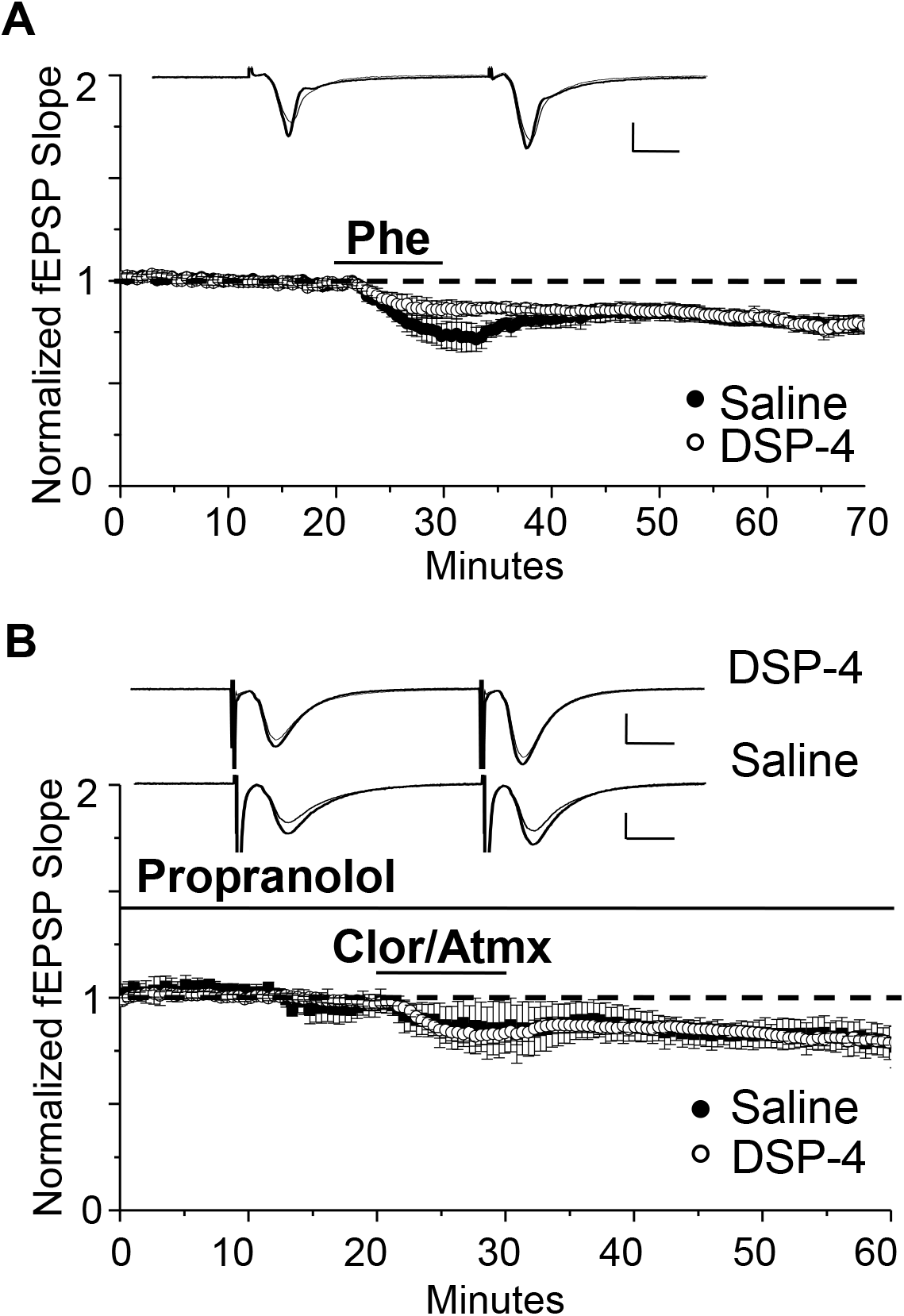
α_1_AR LTD remains intact following NE degeneration and is able to be induced by endogenous NE. A. DSP-4 treatment does not prevent α_1_AR LTD (85±3% of baselined fEPSP slope, n=7). The magnitude of α_1_AR LTD induced by Phe is not significantly different between saline and DSP-4 treatment groups (p >0.05) (control data replotted from Fig 1A). B. α_1_AR LTD is induced by endogenous NE accumulation in slices from DSP-4-treated rats (83±3% of baseline fEPSP slope, n=11). The magnitude of α_1_AR LTD induced by accumulation of endogenous NE is not significantly different between saline and DSP-4 treatment groups (p>0.05) (control data replotted from Fig 2A).

To determine whether the NA fibers surviving neurotoxic damage are able to functionally release NE and activate α_1_ARs effectively to induce LTD, NET, MAO, and β-ARs antagonists were bath applied. Again surprisingly, α_1_AR LTD was observed (Fig. 4B, DSP-4: 83 ± 3%, n=11, p<0.001) and the magnitude was not different from saline-treated animals; (Fig. 4B, saline vs. DSP-4, p=0.97). Furthermore, the magnitude of LTD induced via activation of α_1_ARs by endogenously released NE was not significantly different from that via direct α_1_AR activation by Phe (Figs. 4A and 4B, DSP-4, CAP vs. Phe p=0.58).

To determine if α_1_AR LTD induced by endogenous NE release elicits maximal depression and occludes further depression induced by subsequent application of Phe, the drug mixture CAP was bath applied for 10 minutes to induce LTD. In interleaved slices, Phe (100µM) was coapplied with CAP for 15 minutes beginning 20 minutes after CAP application to determine if additional LTD could be elicited (Fig. 5). We found that no additional LTD could be induced by Phe in the presence of CAP, such that α_1_AR LTD induced by CAP that was not significantly different when Phe was added to the CAP mixture (denoted CAPP) (Fig. 5, CAP: n=6; CAPP: n=10, *t(6)=2.08* p=0.0827).

**Figure 5.**
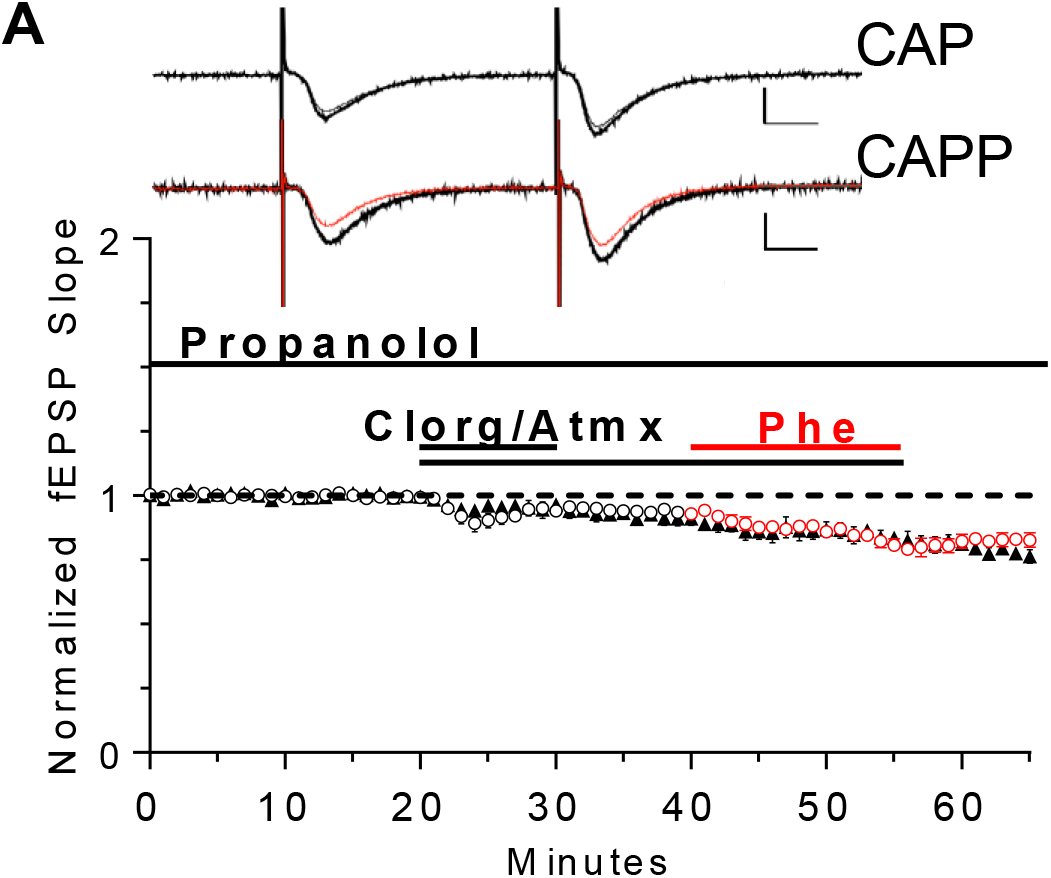
Endogenous NE is sufficient to cause maximal α_1_AR LTD. α_1_AR LTD is induced by endogenous NE accumulation in slices and additional application of Phe is unable to induce further LTD (CAP: n=6; CAPP: n=10, *t(6)=2.08* p=0.0827). Red objects denote the interleaved experiments in which the addition of Phe was included in the CAP mixture following 20 minutes of CAP application.

## Discussion

### α_1_AR LTD

Here we have established that increasing extracellular accumulation of endogenously released NE can activate α_1_ARs and induce α_1_AR LTD at CA3-CA1 synapses in hippocampus. Furthermore, lesion of NA input to hippocampus from the LC did not prevent induction or expression of α_1_AR LTD, despite the loss of ~85% of hippocampal NA innervation. Surprisingly, the NA fibers that remained following DSP-4 induced lesion release enough NE that can accumulate following NET and MAO inhibition to activate postsynaptic α_1_ARs at CA3-CA1 synapses to induce LTD. Therefore, these data suggest that in light of severe NE degeneration, α_1_ ARs remain coupled to their signaling cascade and are able to be activated by specific α_1_AR agonists or via endogenously released NE from surviving NA fibers to induce LTD.

Previously, our lab has shown that M1-acetylcholine receptor (AChR) LTD (mLTD) is lost following degeneration of hippocampal cholinergic innervation from the medial septum, but is rescued when hippocampal NA sympathetic sprouting occurs with an accompanying cholinergic reinnervation of hippocampus at 15% of control levels (Scheiderer et al., 2006). Because M1 mAChRs and α1ARs couple to the same G protein signaling pathway (Porter et al., 2002), and we have shown that their simultaneous weak pharmacological activation induces LTD (Scheiderer et al., 2008), we expected that loss of NA innervation to hippocampus would cause the same deficit in α1AR LTD as we have observed for M1 mAChR LTD following cholinergic degeneration in the absence of sprouting (Scheiderer et al., 2006). However, the successful expression of α_1_ AR LTD at synapses in slices with 15% NA fibers remaining is consistent with the ‘rescued’ mLTD by the approximate 15% cholinergic reinnervation stimulated by NA sympathetic sprouting after medial septal lesion. Thus 15% of control cholinergic or NA innervation in hippocampus is enough to maintain function of M1 mAChRs or α1 ARs, respectively. Based on these findings, we speculate that α_1_ARs remain functional in AD and PD patients even with 67.9% and 83.2% cell loss (Marien et al., 2004). In fact, α_1_ARs expression is increased in AD patients, but unfortunately, this has been associated with increased aggression in these patients (Sharp et al., 2007; Szot et al., 2006). Due to this, prazosin, an α_1_AR antagonist, is being used in AD clinical trials and has been shown to improve aggravation and aggression symptoms (Wang et al., 2010). It is interesting to note that uncoupling of α_1_AR function and loss of α_1_AR LTD occurs at glutamatergic synapses in mPFC in adult rats as a consequence of neonatal lesion of the ventral hippocampus (Bhardwaj et al., 2014).

### 4.2. DSP-4 induced lesion of hippocampal NA innervation

In humans, the severity of LC degeneration is positively correlated with intensity of AD symptoms and is detectable in the prodromal phase, prior to the onset of cognitive deficits (Arendt et al., 2015; Braak et al., 2011; Grudzien et al., 2007). While the onset of this degeneration is driven by tauopathy in humans (Geula et al., 2008; Grinberg et al., 2009; Hertz, 1989; Parvizi et al., 2001; Simic et al., 2009), we were able to recapitulate the effects of AD-driven LC degeneration with the neurotoxin, DSP-4 in healthy rats.

The DSP-4 treatment protocol employed reduced NA innervation by ~85%; additional injections or increases in DSP-4 concentration were not used due to observed increases in animal mortality (unpublished observations). It is important to note that the DSP-4 induced NA lesion is a variable model, as DSP-4 only provides a temporary decrease in NA innervation, whereas neurodegeneration is permanent. In fact, several studies have shown that DSP-4 induced NA degeneration is reversed several months following treatment (Fritschy et al., 1990; Fritschy and Grzanna, 1989, 1992). The LC-NA system is known to be capable of initiating these compensatory mechanisms in response to damage, which includes an increase in NE turnover (Jonsson et al., 1981) and release (Abercrombie and Zigmond, 1989) in surviving cell axons, as well as inducing α and β receptor supersensitivity (Berridge and Dunn, 1990; Dooley et al., 1983; Starke, 2001). Interestingly, AD patients have elevated NE levels even though there is LC cell loss (Szot et al., 2006), and α_2_-AR positive axonal sprouting has been identified in AD post-mortem human hippocampus (Szot et al., 2006). Thus, it can be postulated that excessive release, in addition to depletion, of NE could also lead to deficits in hippocampal dependent learning and memory, and therefore, synaptic plasticity. Furthermore, the lesion protocol used here may cause increased activation of the NA compensatory mechanisms thought to be responsible for lesion reversal (Fritschy and Grzanna, 1992). In support of this idea, the DβH positive fiber morphology appears thicker and has a knobby appearance in the DSP-4 treated group compared to saline-treated control (Fig. 3A arrows). This morphology change could be an early indication of compensatory LC sprouting because as mentioned above, it has been shown DSP-4 induced lesions are not permanent, and after a variable period of time hyperinnervation of NA fibers can occur (Booze et al., 1988; Fritschy and Grzanna, 1992; Kalinin et al., 2006). Altogether, we were unable to induce a complete loss of all NA hippocampal innervation using DSP-4 and were therefore unable to observe α_1_AR uncoupling and α_1_AR LTD loss that we predict would happen in the complete absence of endogenous NE, similar to our previous studies of M1mAChR LTD following complete medial septal lesion (Scheiderer et al., 2006).

The maintenance of α_1_AR function we observe following DSP-4 induce lesion could explain the inconsistent findings in learning and memory assays following DSP-4 treatment, with some studies reporting minimal effects (Decker and McGaugh, 1991; Ohno et al., 1993, 1997; Prado de Carvalho and Zornetzer, 1981), whereas others find deficits (Decker and McGaugh, 1991; Prado de Carvalho and Zornetzer, 1981). The reversibility of DSP-4 treatment, as well as variable treatment paradigms could lead to the confounding behavioral results following NA degeneration.

## Conclusions

Here we show that pharmacological inhibition of NET and MAO leads to extracellular accumulation of NE which is capable of activating ARs that modulate synaptic efficacy at CA3-CA1 synapses. This finding is important, as NET and MAO inhibitors are used clinically in the treatment of several neurological and neuropsychiatric illness (Castellanos et al., 1996; Zametkin and Rapoport, 1987). Furthermore, the DSP-4-driven loss of ~85% of NA fibers in hippocampus recapitulates the loss cortical projecting LC-cells in human AD (Arendt et al., 2015; Braak et al., 2011; German et al., 1992; Grudzien et al., 2007). The present findings suggest that α_1_ARs are preserved in CA3-CA1 glutamatergic synapses following extensive NA fiber denervation. This preservation of α_1_AR function may be an effect of compensatory NE levels released from remaining fibers or may be resilient to change in chronically low stimulation settings. These results indicate that NA drugs may provide an avenue for viable therapeutics in AD to treat changes in local NE levels in regions critical for learning and memory.

## Author Contributions

KDR and LLM conceived and designed the study; KDR and AMG contributed to the execution of the experiments and performed the statistical analyses; KDR wrote the first draft of the manuscript; ARN, AMG, and LLM wrote sections of the manuscript. All authors contributed to the manuscript revision, read and approved the submitted version

## Acknowledgments

We would like to thank the UAB-High Resolution Imaging Facility for use of their confocal microscope.

## Data Availability Statement

Datasets are available on request:

The raw data supporting the conclusions of this manuscript will be made available by the authors, without undue reservation, to any qualified researcher.

## References

Abercrombie, E. D., and Zigmond, M. J. (1989). Partial injury to central noradrenergic neurons: reduction of tissue norepinephrine content is greater than reduction of extracellular norepinephrine measured by microdialysis. J. Neurosci. 9, 4062–4067.

André, M. A. E., Wolf, O. T., and Manahan-Vaughan, D. (2015). Beta-adrenergic receptors support attention to extinction learning that occurs in the absence, but not the presence, of a context change. Front. Behav. Neurosci. 9, 125. doi:10.3389/fnbeh.2015.00125.

Arendt, T., Bruckner, M. K., Morawski, M., Jager, C., and Gertz, H. J. (2015). Early neurone loss in Alzheimer’s disease: cortical or subcortical? Acta Neuropathol Commun 3, 10. doi:10.1186/s40478-015-0187-1.

Aston-Jones, G. (2004). “CHAPTER 11 – Locus Coeruleus, A5 and A7 Noradrenergic Cell Groups,” in The Rat Nervous System, 259–294. doi:10.1016/B978-012547638-6/50012-2.

Berridge, C. W., and Dunn, A. J. (1990). DSP-4-induced depletion of brain norepinephrine produces opposite effects on exploratory behavior 3 and 14 days after treatment. Psychopharmacology (Berl). 100, 504–8. Available at: http://www.ncbi.nlm.nih.gov/pubmed/2320711.

Bhardwaj, S. K., Tse, Y. C. hun., Ryan, R., Wong, T. P. a., and Srivastava, L. K. (2014). Impaired adrenergic-mediated plasticity of prefrontal cortical glutamate synapses in rats with developmental disruption of the ventral hippocampus. Neuropsychopharmacology 39, 2963–2973. doi:10.1038/npp.2014.142.

Booze, R. M., Hall, J. A., Cress, N. M., Miller, G. D., and Davis, J. N. (1988). DSP-4 treatment produces abnormal tyrosine hydroxylase immunoreactive fibers in rat hippocampus. Exp. Neurol. 101, 75–86. doi:10.1016/0014-4886(88)90066-0.

Braak, H., Thal, D. R., Ghebremedhin, E., and Del Tredici, K. (2011). Stages of the Pathologic Process in Alzheimer Disease: Age Categories From 1 to 100 Years. J. Neuropathol. Exp. Neurol. 70, 960–969. doi:10.1097/NEN.0b013e318232a379.

Bramham, C. R., Bacher-Svendsen, K., and Sarvey, J. M. (1997). LTP in the lateral perforant path is beta-adrenergic receptor-dependent. Neuroreport 8, 719–24. Available at: http://insights.ovid.com/pubmed?pmid=9106754.

Bröcher, S., Artola, A., and Singer, W. (1992). Agonists of cholinergic and noradrenergic receptors facilitate synergistically the induction of long-term potentiation in slices of rat visual cortex. Brain Res. 573, 27–36. doi:10.1016/0006-8993(92)90110-U.

Castellanos, F. X., Giedd, J. N., Marsh, W. L., Hamburger, S. D., Vaituzis, A. C., Dickstein, D. P., et al. (1996). Quantitative brain magnetic resonance imaging in attention-deficit hyperactivity disorder. Arch. Gen. Psychiatry 53, 607–16. Available at: http://www.ncbi.nlm.nih.gov/pubmed/8660127.

Chalermpalanupap, T., Weinshenker, D., and Rorabaugh, J. M. (2017). Down but Not Out: The Consequences of Pretangle Tau in the Locus Coeruleus. Neural Plast. 2017. doi:10.1155/2017/7829507.

Collette, K. M., Zhou, X. D., Amoth, H. M., Lyons, M. J., Papay, R. S., Sens, D. A., et al. (2014). Long-term α1B-adrenergic receptor activation shortens lifespan, while α1A-adrenergic receptor stimulation prolongs lifespan in association with decreased cancer incidence. Age (Omaha). 36. doi:10.1007/s11357-014-9675-7.

Decker, M. W., and McGaugh, J. L. (1991). The role of interactions between the cholinergic system and other neuromodulatory systems in learning and memory. Synapse 7, 151–68. doi:10.1002/syn.890070209.

Dooley, D. J., Mogilnicka, E., Delini-Stula, A., Waechter, F., Truog, A., and Wood, J. (1983). Functional supersensitivity to adrenergic agonists in the rat after DSP-4, a selective noradrenergic neurotoxin. Psychopharmacology (Berl). 81, 1–5. doi:10.1007/BF00439263.

Doze, V. a, Papay, R. S., Goldenstein, B. L., Gupta, M. K., Collette, K. M., Nelson, B. W., et al. (2011). Long-Term alpha-1A Adrenergic Receptor Stimulation Improves Synaptic Plasticity, Cognitive Function, Mood, and Longevity. Mol. Pharmacol. 80, 747–758. doi:10.1124/mol.111.073734.Norepinephrine.

Erickson, J. C., Hollopeter, G., Thomas, S. A., Froelick, G. J., and Palmiter, R. D. (1997). Disruption of the metallothionein-III gene in mice: analysis of brain zinc, behavior, and neuron vulnerability to metals, aging, and seizures. J. Neurosci. 17, 1271–1281.

Forno, L. S. (1966). Pathology of parkinsonism-A preliminary report of 24 cases. J. Neurosurg. 24, 266.

Fritschy, J. M., Geffard, M., and Grzanna, R. (1990). The response of noradrenergic axons to systemically administered DSP-4 in the rat: an immunohistochemical study using antibodies to noradrenaline and dopamine-beta-hydroxylase. J. Chem. Neuroanat. 3, 309–21. Available at: http://www.ncbi.nlm.nih.gov/pubmed/2204356.

Fritschy, J. M., and Grzanna, R. (1989). Immunohistochemical analysis of the neurotoxic effects of DSP-4 identifies two populations of noradrenergic axon terminals. Neuroscience 30, 181–197. doi:10.1016/0306-4522(89)90364-3.

Fritschy, J. M., and Grzanna, R. (1992). Restoration of ascending noradrenergic projections by residual locus coeruleus neurons: compensatory response to neurotoxin-induced cell death in the adult rat brain. J. Comp. Neurol. 321, 421–41. doi:10.1002/cne.903210309.

German, D., Manaye, K., White, C., Woodward, D., McIntire, D., Smith, W., et al. (1992). Disease-specific patterns of locus coeruleus cell loss. Annu. Neurol. 32, 667–676. doi:10.1002/ana.410320510.

Geula, C., Nagykery, N., Nicholas, A., and Wu, C. (2008). Cholinergic neuronal and axonal abnormalities are present early in aging and in Alzheimer disease. J. Neuropathol. Exp. Neurol. 67, 309–18. doi:10.1097/NEN.0b013e31816a1df3.

Gibbs, M. E., Hutchinson, D. S., and Summers, R. J. (2010). Noradrenaline release in the locus coeruleus modulates memory formation and consolidation; roles for α- and β-adrenergic receptors. Neuroscience 170, 1209–1222. doi:10.1016/j.neuroscience.2010.07.052.

Grinberg, L. T., Rüb, U., Ferretti, R. E. L., Nitrini, R., Farfel, J. M., Polichiso, L., et al. (2009). The dorsal raphe nucleus shows phospho-tau neurofibrillary changes before the transentorhinal region in Alzheimer̈s disease. A precocious onset? Neuropathol. Appl. Neurobiol. 35, 405–416. doi:10.1111/j.1365-2990.2008.00997.x.

Grudzien, A., Shaw, P., Weintraub, S., Bigio, E., Mash, D. C., and Mesulam, M. M. (2007). Locus coeruleus neurofibrillary degeneration in aging, mild cognitive impairment and early Alzheimer’s disease. Neurobiol. Aging 28, 327–335. doi:10.1016/j.neurobiolaging.2006.02.007.

Hagena, H., Hansen, N., and Manahan-Vaughan, D. (2016). β-Adrenergic Control of Hippocampal Function: Subserving the Choreography of Synaptic Information Storage and Memory. Cereb. Cortex 26, 1349–1364. doi:10.1093/cercor/bhv330.

Harley, C. (1991). Noradrenergic and locus coeruleus modulation of the perforant path-evoked potential in rat dentate gyrus supports a role for the locus coeruleus in attentional and memorial processes. Elsevier B.V. doi:10.1016/S0079-6123(08)63818-2.

Harley, C. W. (2007). Norepinephrine and the dentate gyrus. Prog. Brain Res. 163, 299–318. doi:10.1016/S0079-6123(07)63018-0.

Harley, C. W., and Sara, S. J. (1992). Locus coeruleus bursts induced by glutamate trigger delayed perforant path spike amplitude potentiation in the dentate gyrus. Exp. Brain Res. 89, 581–587. doi:10.1007/BF00229883.

Harro, J., Pähkla, R., Modiri, A. R., Harro, M., Kask, A., and Oreland, L. (1999). Dose-dependent effects of noradrenergic denervation by DSP-4 treatment on forced swimming and β-adrenoceptor binding in the rat. J. Neural Transm. 106, 619–629. doi:10.1007/s007020050184.

Hertz, L. (1989). Is Alzheimer’s disease an anterograde degeneration, originating in the brainstem, and disrupting metabolic and functional interactions between neurons and glial cells? Brain Res. Rev. 14, 335–353. doi:10.1016/0165-0173(89)90017-9.

Hopkins, W. F., and Johnston, D. (1984). Frequency-dependent noradrenergic modulation of long-term potentiation in the hippocampus. Science 226, 350–2. Available at: http://www.ncbi.nlm.nih.gov/pubmed/6091272.

Huang, Y. Y., Nguyen, P. V, Abel, T., and Kandel, E. R. (1996). Long-lasting forms of synaptic potentiation in the mammalian hippocampus. Learn. Mem. 3, 74–85. doi:10.1101/lm.3.2-3.74.

Israel, J. A. (2015). Combining Stimulants and Monoamine Oxidase Inhibitors: A Reexamination of the Literature and a Report of a New Treatment Combination. Prim. care companion CNS Disord. 17. doi:10.4088/PCC.15br01836.

Izumi, Y., and Zorumski, C. F. (1999). Norepinephrine promotes long-term potentiation in the adult rat hippocampus in vitro. Synapse 31, 196–202. doi:10.1002/(SICI)1098-2396(19990301)31:3<196::AID-SYN4>3.0.CO;2-K.

Jaim-Etcheverry, G., and Zieher, L. M. (1980). DSP-4: a novel compound with neurotoxic effects on noradrenergic neurons of adult and developing rats. Brain Res. 188, 513–523. doi:10.1016/0006-8993(80)90049-9.

Janitzky, K., Lippert, M. T., Engelhorn, A., Tegtmeier, J., Goldschmidt, J., Heinze, H.-J., et al. (2015). Optogenetic silencing of locus coeruleus activity in mice impairs cognitive flexibility in an attentional set-shifting task. Front. Behav. Neurosci. 9. doi:10.3389/fnbeh.2015.00286.

Jonsson, G., Hallman, H., Ponzio, F., and Ross, S. (1981). DSP4 (N-(2-chloroethyl)-N-ethyl-2-bromobenzylamine)-A useful denervation tool for central and peripheral noradrenaline neurons. Eur. J. Pharmacol. 72, 173–188. doi:10.1016/0014-2999(81)90272-7.

Jucker, M., and Walker, L. C. (2011). Pathogenic protein seeding in Alzheimer disease and other neurodegenerative disorders. Ann. Neurol. 70, 532–40. doi:10.1002/ana.22615.

Kalinin, S., Feinstein, D. L., Xu, H. L., Huesa, G., Pelligrino, D. A., and Galea, E. (2006). Degeneration of noradrenergic fibres from the locus coeruleus causes tight-junction disorganisation in the rat brain. Eur. J. Neurosci. 24, 3393–3400. doi:10.1111/j.1460-9568.2006.05223.x.

Katsuki, H., Izumi, Y., and Zorumski, C. F. (1997). Noradrenergic regulation of synaptic plasticity in the hippocampal CA1 region. J. Neurophysiol. 77, 3013–20. Available at: http://www.ncbi.nlm.nih.gov/pubmed/9212253.

Kelly, S. C., He, B., Perez, S. E., Ginsberg, S. D., Mufson, E. J., and Counts, S. E. (2017). Locus coeruleus cellular and molecular pathology during the progression of Alzheimer’s disease. Acta Neuropathol. Commun. 5, 8. doi:10.1186/s40478-017-0411-2.

Kemp, A., and Manahan-Vaughan, D. (2008). β-Adrenoreceptors comprise a critical element in learning-facilitated long-term plasticity. Cereb. Cortex 18, 1326–1334. doi:10.1093/cercor/bhm164.

Koob, G. F., Kelley, A. E., and Mason, S. T. (1978). Locus coeruleus lesions: Learning and extinction. Physiol. Behav. 20, 709–716. doi:10.1016/0031-9384(78)90296-2.

Lemon, N., Aydin-Abidin, S., Funke, K., and Manahan-Vaughan, D. (2009). Locus coeruleus activation facilitates memory encoding and induces hippocampal LTD that depends on β-Adrenergic receptor activation. Cereb. Cortex 19, 2827–2837. doi:10.1093/cercor/bhp065.

Mann, D. M. A. (1983). The locus coeruleus and its possible role in ageing and degenerative disease of the human central nervous system. Mech. Ageing Dev. 23, 73–94. doi:10.1016/0047-6374(83)90100-8.

Mann, D. M. A., Yates, P. O., and Hawkes, J. (1983). The pathology of the human locus ceruleus. Clin. Neuropathol. 2, 1–7. Available at: http://www.ncbi.nlm.nih.gov/pubmed/6220852.

Marien, M. R., Colpaert, F. C., and Rosenquist, A. C. (2004). Noradrenergic mechanisms in neurodegenerative diseases: A theory. Brain Res. Rev. 45, 38–78. doi:10.1016/j.brainresrev.2004.02.002.

Ohno, M., Yamamoto, T., Kobayashi, M., and Watanabe, S. (1993). Impairment of working memory induced by scopolamine in rats with noradrenergic DSP-4 lesions. Eur. J. Pharmacol. 238, 117–120. doi:10.1016/0014-2999(93)90514-I.

Ohno, M., Yoshimatsu, A., Kobayashi, M., and Watanabe, S. (1997). Noradrenergic DSP-4 lesions aggravate impairment of working memory produced by hippocampal muscarinic blockade in rats. Pharmacol. Biochem. Behav. 57, 257–261. doi:10.1016/S0091-3057(96)00353-X.

Parvizi, J., Van Hoesen, G. W., and Damasio, A. (2001). The selective vulnerability of brainstem nuclei to Alzheimer’s disease. Ann. Neurol. 49, 53–66. doi:10.1002/1531-8249(200101)49:1<53::AID-ANA30>3.0.CO;2-Q.

Porter, A. C., Bymaster, F. P., DeLapp, N. W., Yamada, M., Wess, J., Hamilton, S. E., et al. (2002). M1 muscarinic receptor signaling in mouse hippocampus and cortex. Brain Res. 944, 82–89. doi:10.1016/S0006-8993(02)02721-X.

Prado de Carvalho, L., and Zornetzer, S. F. (1981). The involvement of the locus coeruleus in memory. Behav. Neural Biol. 31, 173–186. doi:10.1016/S0163-1047(81)91204-8.

Puoliväli, J., Pradier, L., and Riekkinen, P. (2000). Impaired recovery of noradrenaline levels in apolipoprotein E-deficient mice after N-(2-chloroethyl)-N-ethyl-2-bromobenzylamine lesion. Neuroscience 95, 353–8. Available at: http://www.ncbi.nlm.nih.gov/pubmed/10658614.

Pussinen, R., Nieminen, S., Koivisto, E., Haapalinna, a, Riekkinen, P., and Sirvio, J. (1997). Enhancement of intermediate-term memory by an alpha-1 agonist or a partial agonist at the glycine site of the NMDA receptor. Neurobiol. Learn. Mem. 67, 69–74. doi:10.1006/nlme.1996.3738.

Puumala, T., Greijus, S., Narinen, K., Haapalinna, A., Riekkinen, P., and Sirviö, J. (1998). Stimulation of alpha-1 adrenergic receptors facilitates spatial learning in rats. Eur. Neuropsychopharmacol. 8, 17–26. doi:10.1016/S0924-977X(97)00040-0.

Ross, S. B. (1976). Long-term effects of N-2-chlorethyl-N-ethyl-2-bromobenzylamine hydrochloride on noradrenergic neurones in the rat brain and heart. Br. J. Pharmacol. 58, 521–7. Available at: http://www.ncbi.nlm.nih.gov/pubmed/1000130%5Cnhttp://www.pubmedcentral.nih.gov/articlerender.fcgi?artid=PMC1667482.

Ross, S. B., and Stenfors, C. (2014). DSP4, a Selective Neurotoxin for the Locus Coeruleus Noradrenergic System. A Review of Its Mode of Action. Neurotox. Res., 15–30. doi:10.1007/s12640-014-9482-z.

Scheiderer, C. L., Dobrunz, L. E., and McMahon, L. L. (2004). Novel form of long-term synaptic depression in rat hippocampus induced by activation of α1 adrenergic receptors. J. Neurophysiol. 91, 1071–1077. doi:10.1152/jn.00420.2003.

Scheiderer, C. L., McCutchen, E., Thacker, E. E., Kolasa, K., Ward, M. K., Parsons, D., et al. (2006). Sympathetic sprouting drives hippocampal cholinergic reinnervation that prevents loss of a muscarinic receptor-dependent long-term depression at CA3-CA1 synapses. J. Neurosci. 26, 3745–56. doi:10.1523/JNEUROSCI.5507-05.2006.

Scheiderer, C. L., Smith, C. C., McCutchen, E., McCoy, P. A., Thacker, E. E., Kolasa, K., et al. (2008). Coactivation of M1 Muscarinic and 1 Adrenergic Receptors Stimulates Extracellular Signal-Regulated Protein Kinase and Induces Long-Term Depression at CA3-CA1 Synapses in Rat Hippocampus. J. Neurosci. 28, 5350–5358. doi:10.1523/JNEUROSCI.5058-06.2008.

Sharp, S. I., Ballard, C. G., Chen, C. P. L.-H., and Francis, P. T. (2007). Aggressive behavior and neuroleptic medication are associated with increased number of alpha1-adrenoceptors in patients with Alzheimer disease. Am. J. Geriatr. Psychiatry 15, 435–7. doi:10.1097/01.JGP.0000237065.78966.1b.

Simic, G., Stanic, G., Mladinov, M., Jovanov-Milosevic, N., Kostovic, I., and Hof, P. R. (2009). Does Alzheimer’s disease begin in the brainstem?: Annotation. Neuropathol. Appl. Neurobiol. 35, 532–554. doi:10.1111/j.1365-2990.2009.01038.x.

Starke, K. (2001). Presynaptic autoreceptors in the third decade: Focus on ??2-adrenoceptors. J. Neurochem. 78, 685–693. doi:10.1046/j.1471-4159.2001.00484.x.

Szot, P. (2012). Common factors among Alzheimer’s disease, Parkinson’s disease, and epilepsy: Possible role of the noradrenergic nervous system. Epilepsia 53, 61–66. doi:10.1111/j.1528-1167.2012.03476.x.

Szot, P., White, S. S., Greenup, J. L., Leverenz, J. B., Peskind, E. R., and Raskind, M. A. (2006). Compensatory changes in the noradrenergic nervous system in the locus ceruleus and hippocampus of postmortem subjects with Alzheimer’s disease and dementia with Lewy bodies. J. Neurosci. 26, 467–78. doi:10.1523/JNEUROSCI.4265-05.2006.

Theofilas, P., Ehrenberg, A. J., Dunlop, S., Di Lorenzo Alho, A. T., Nguy, A., Leite, R. E. P., et al. (2017). Locus coeruleus volume and cell population changes during Alzheimer’s disease progression: A stereological study in human postmortem brains with potential implication for early-stage biomarker discovery. Alzheimer’s Dement. 13, 236–246. doi:10.1016/j.jalz.2016.06.2362.

Thomas, S. A., and Palmiter, R. D. (1997a). Disruption of the dopamine Β-hydroxylase gene in mice suggests roles for norepinephrine in motor function, learning, and memory. Behav. Neurosci. 111, 579–589. doi:10.1037/0735-7044.111.3.579.

Thomas, S. A., and Palmiter, R. D. (1997b). Impaired maternal behavior in mice lacking norepinephrine and epinephrine. Cell 91, 583–592. doi:10.1016/S0092-8674(00)80446-8.

Thomas, S. A., and Palmiter, R. D. (1997c). Thermoregulatory and metabolic phenotypes of mice lacking noradrenaline and adrenaline. Nature 387, 94–97. doi:10.1038/387094a0.

Vanicek, T., Spies, M., Rami-Mark, C., Savli, M., Höflich, A., Kranz, G. S., et al. (2014). The norepinephrine transporter in attention-deficit/hyperactivity disorder investigated with positron emission tomography. JAMA psychiatry 71, 1340–1349. doi:10.1001/jamapsychiatry.2014.1226.

Wang, L. Y., Shofer, J. B., Rohde, K., Hart, K. L., Hoff, D. J., McFall, Y. H., et al. (2010). Prazosin for the Treatment of Behavioral Symptoms in Alzheimer’S Disease Patients With Agitation and Aggression. Am. J. Geriatr. Psych 17, 744–751. doi:10.1097/JGP.0b013e3181ab8c61.PRAZOSIN.

Yamada, M., and Mehraein, P. (1977). Verteilungsmuster der senilen Veränderungen in den Hirnstammkernen. Psychiatry Clin. Neurosci. 31, 219–224. doi:10.1111/j.1440-1819.1977.tb02722.x.

Youdim, M. B., and Riederer, P. (1993). Dopamine metabolism and neurotransmission in primate brain in relationship to monoamine oxidase A and B inhibition. J. Neural Transm. Gen. Sect. 91, 181–95. Available at: http://www.ncbi.nlm.nih.gov/pubmed/8390270.

Zametkin, A. J., and Rapoport, J. L. (1987). Neurobiology of Attention Deficit Disorder With Hyperactivity: Where Have We Come in 50 Years? J. Am. Acad. Child Adolesc. Psychiatr. 26, 676–686. doi:10.1097/00004583-198709000-00011.

Zarow, C., Lyness, S. a, Mortimer, J. a, and Chui, H. C. (2003). Neuronal loss is greater in the locus coeruleus than nucleus basalis and substantia nigra in Alzheimer and Parkinson diseases. Arch. Neurol. 60, 337–341. doi:10.1001/archneur.60.3.337.

